## Supplemental Information for "Structural evolution of nitrogenase enzymes over geologic time"

**TITLE**

**SHORT TITLE**

Three billion years of nitrogenase diversity

**AUTHORS AND AFFILIATIONS**

Bruno Cuevas-Zuviría^1,2†^, Franka Detemple^3^, Kaustubh Amritkar^1^, Amanda K. Garcia, Lance Seefeldt^4^, Oliver Einsle^3^ and Betül Kaçar^1*^

1. Department of Bacteriology, University of Wisconsin-Madison, Madison, United States
2. Centro de Biotecnología y Genómica de Plantas (UPM-INIA/CSIC), Universidad Politécnica de Madrid (UPM)—Instituto Nacional de Investigación y Tecnología Agraria y Alimentaria-CSIC (INIA/CSIC), Campus de Montegancedo, Madrid, Spain
3. Institute of Biochemistry, University of Freiburg, Freiburg, Germany
4. Department of Chemistry and Biochemistry, Utah State University, Logan, United States

^†^Affiliation where research was conducted: Department of Bacteriology, University of Wisconsin-Madison, Madison, United States

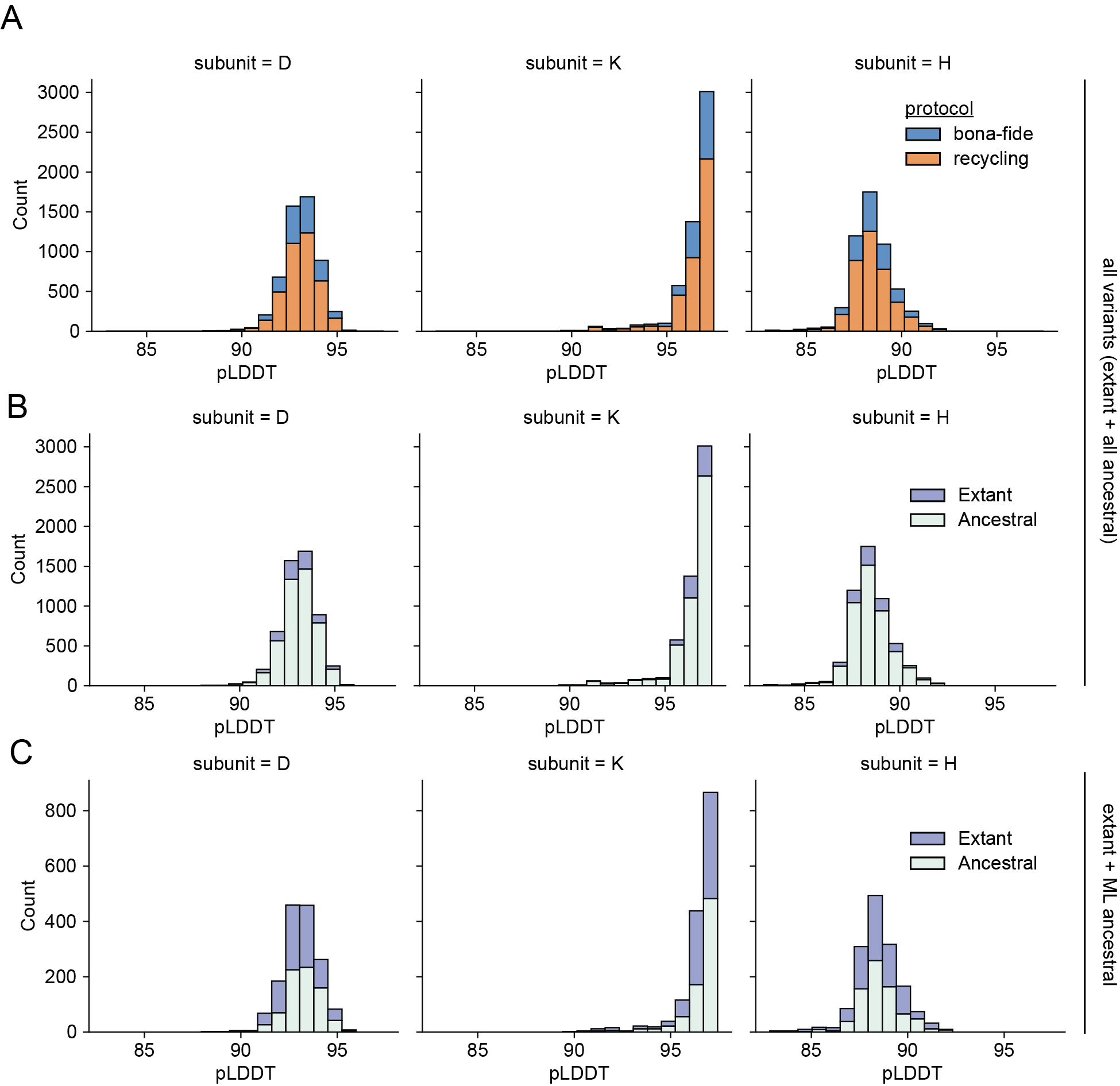

**Figure S1. AlphaFold prediction confidence for extant and ancestral nitrogenases. A)** Histograms plot counts of individual chain predictions (5378 total structures (DDKK + HH) for 769 tree nodes, including six alternative ancestors for each ancestral node), demonstrated for each nitrogenase subunit, colored by the protocol that was employed to produce the structures. **B)** Same data as in panel A, but structures are colored based on whether they represent extant or ancestral nitrogenases. **C)** Histograms of individual chain predictions specifically for extant nitrogenases and their most likely ancestors. This subset includes 1,538 total structures (DDKK + HH) corresponding to the same 769 tree nodes.

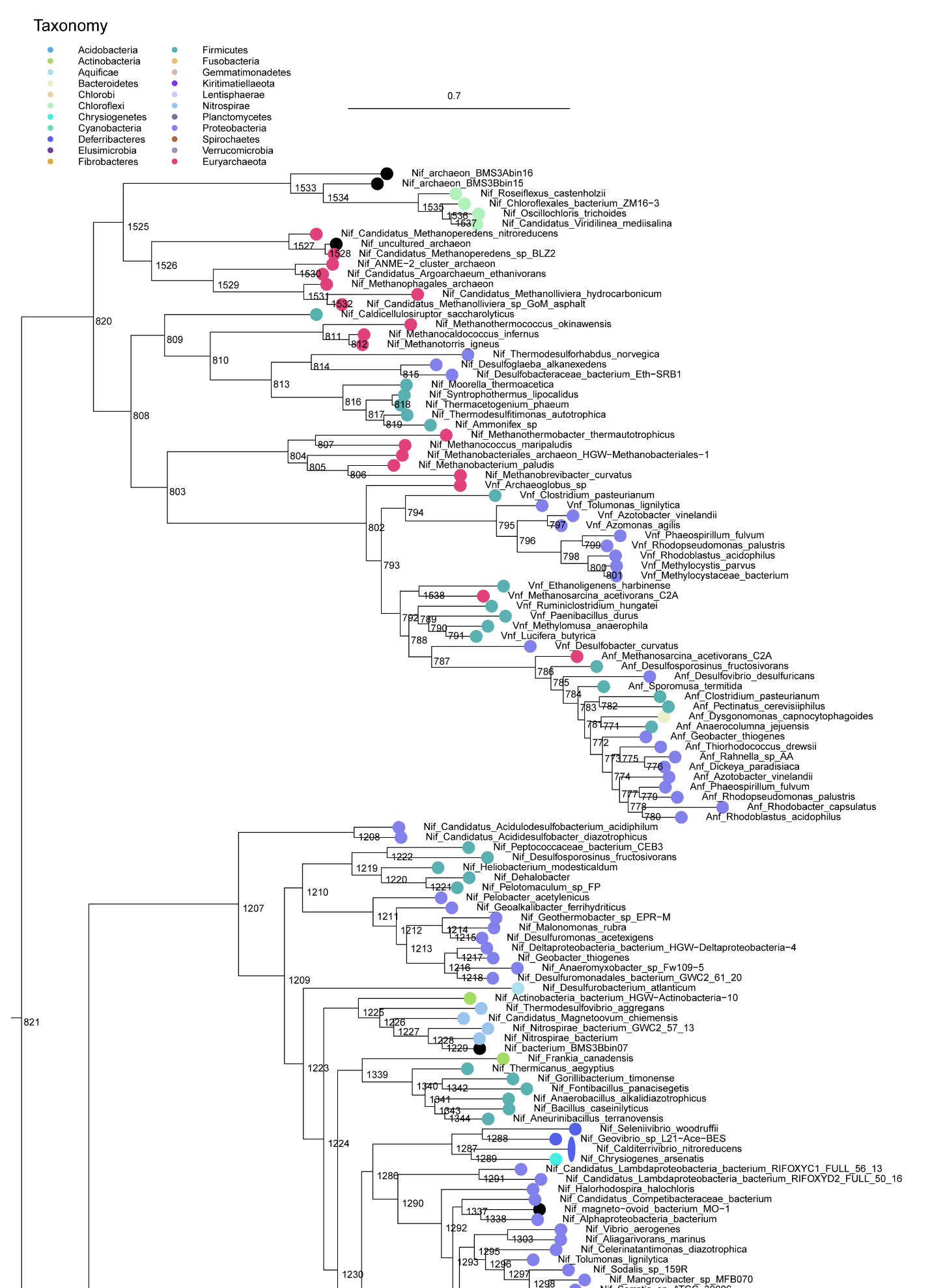

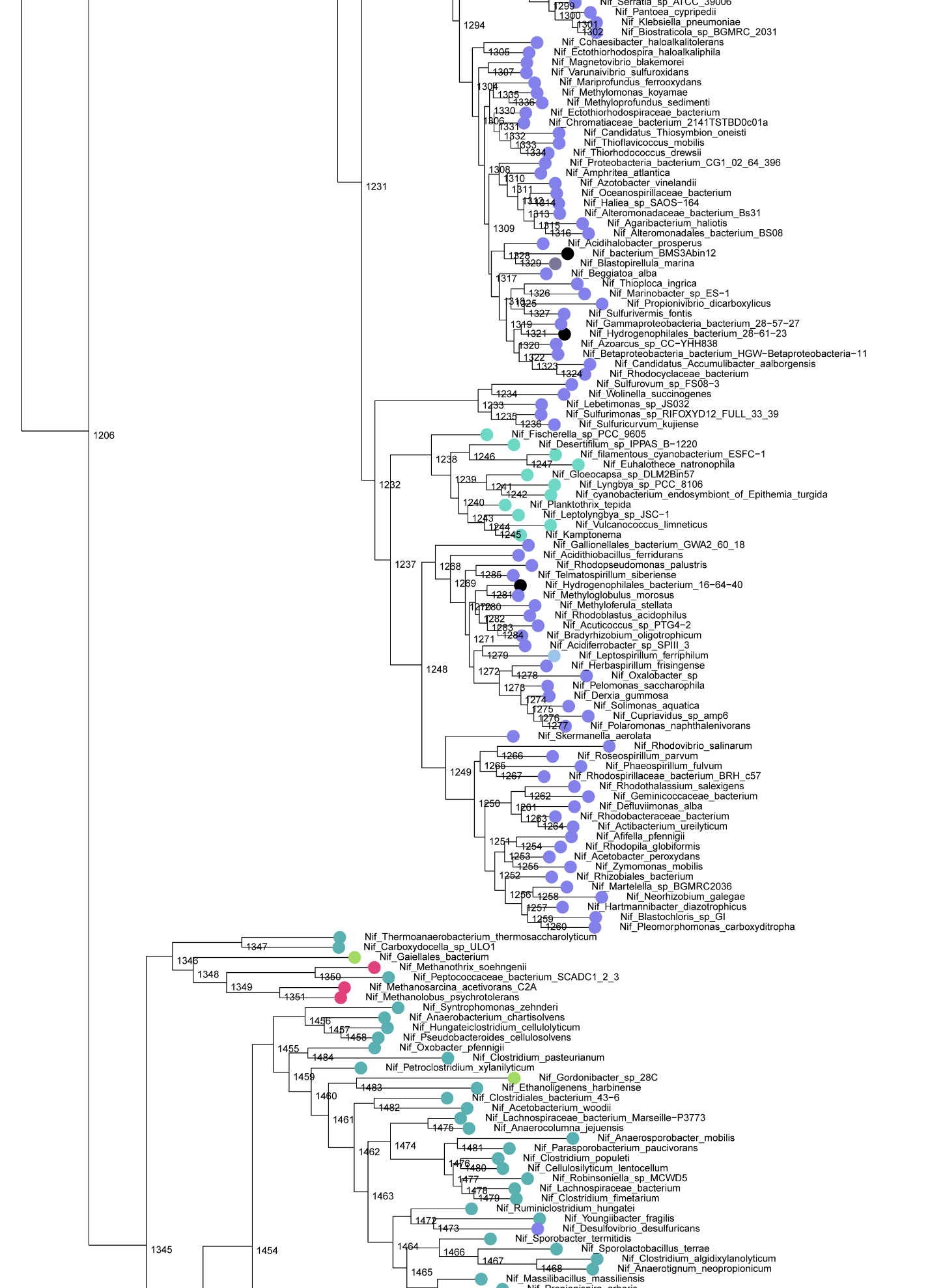

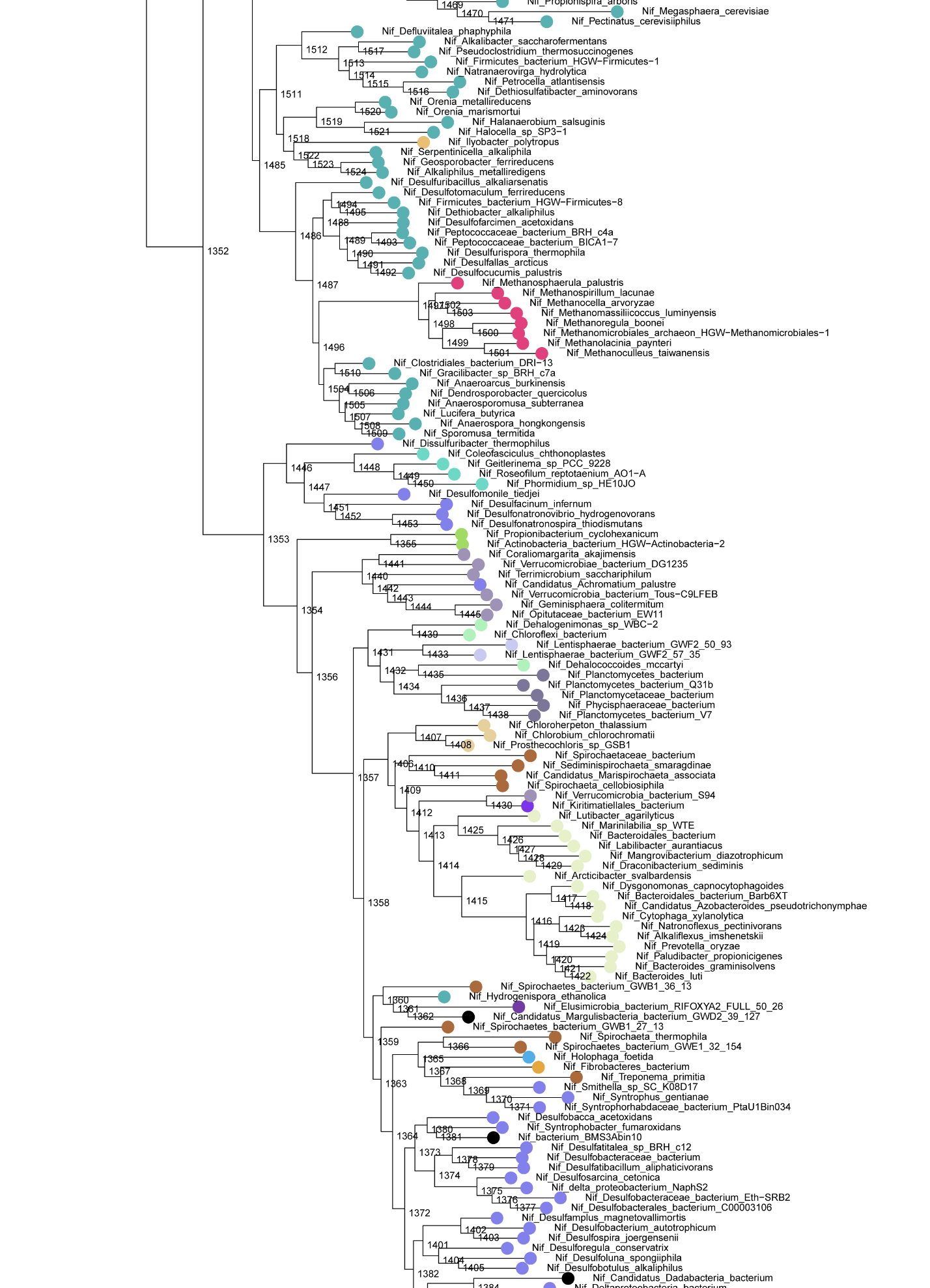

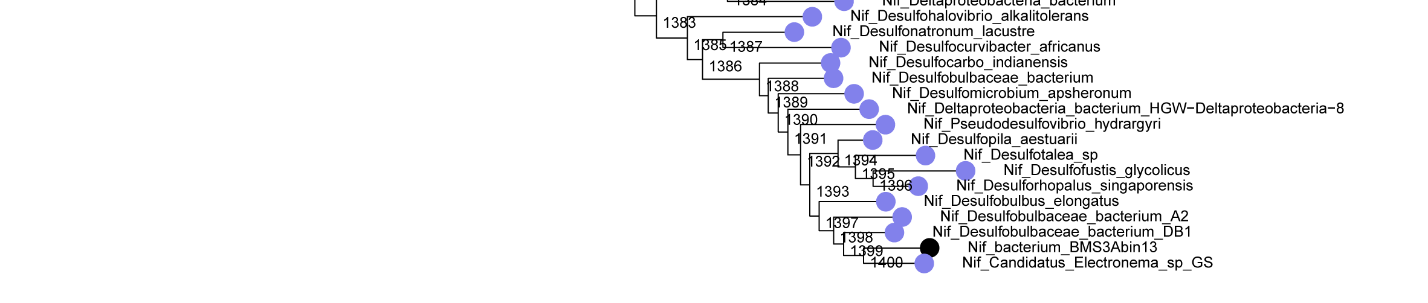

**Figure S2. Expanded n­­itrogenase protein sequence phylogeny** (adapted from Garcia et al., 2020).

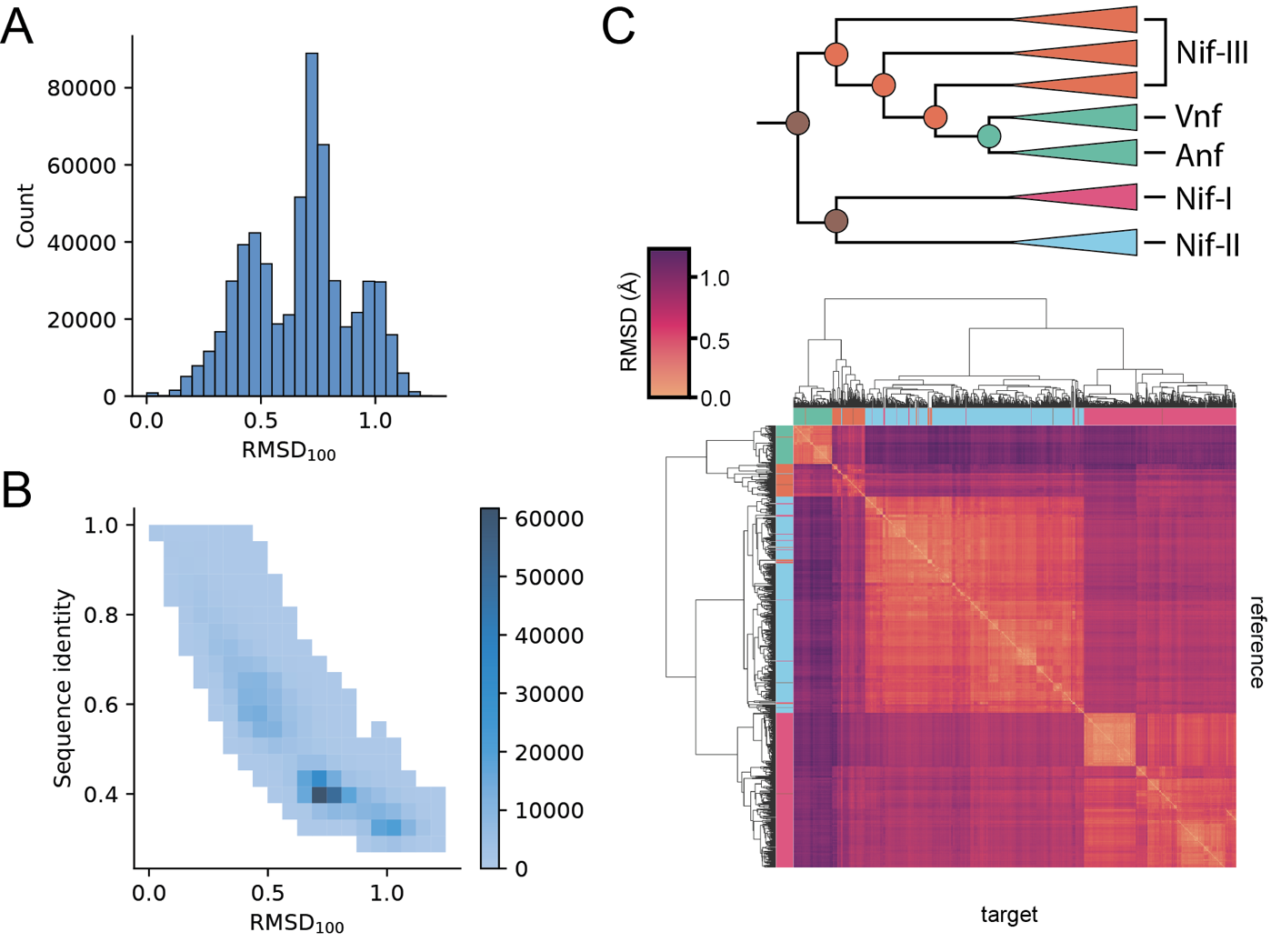

**Figure S3**: **DDKK RMSD_100_ relationships. A)** RMSD_100_ histogram. RMSD_100_ consists on a normalization of the RMSD values shown in **Figure 2** multiplied by a factor that accounts for alignment length. **B)** Bi-dimensional histogram relating RMSD_100_ and sequence identity. **C)** Hierarchical clustering of predicted nitrogenase structures based on structural similarity (RMSD_100_). The colors on the heatmap correspond to the phylogenetic groups represented on **Figure 2C**.

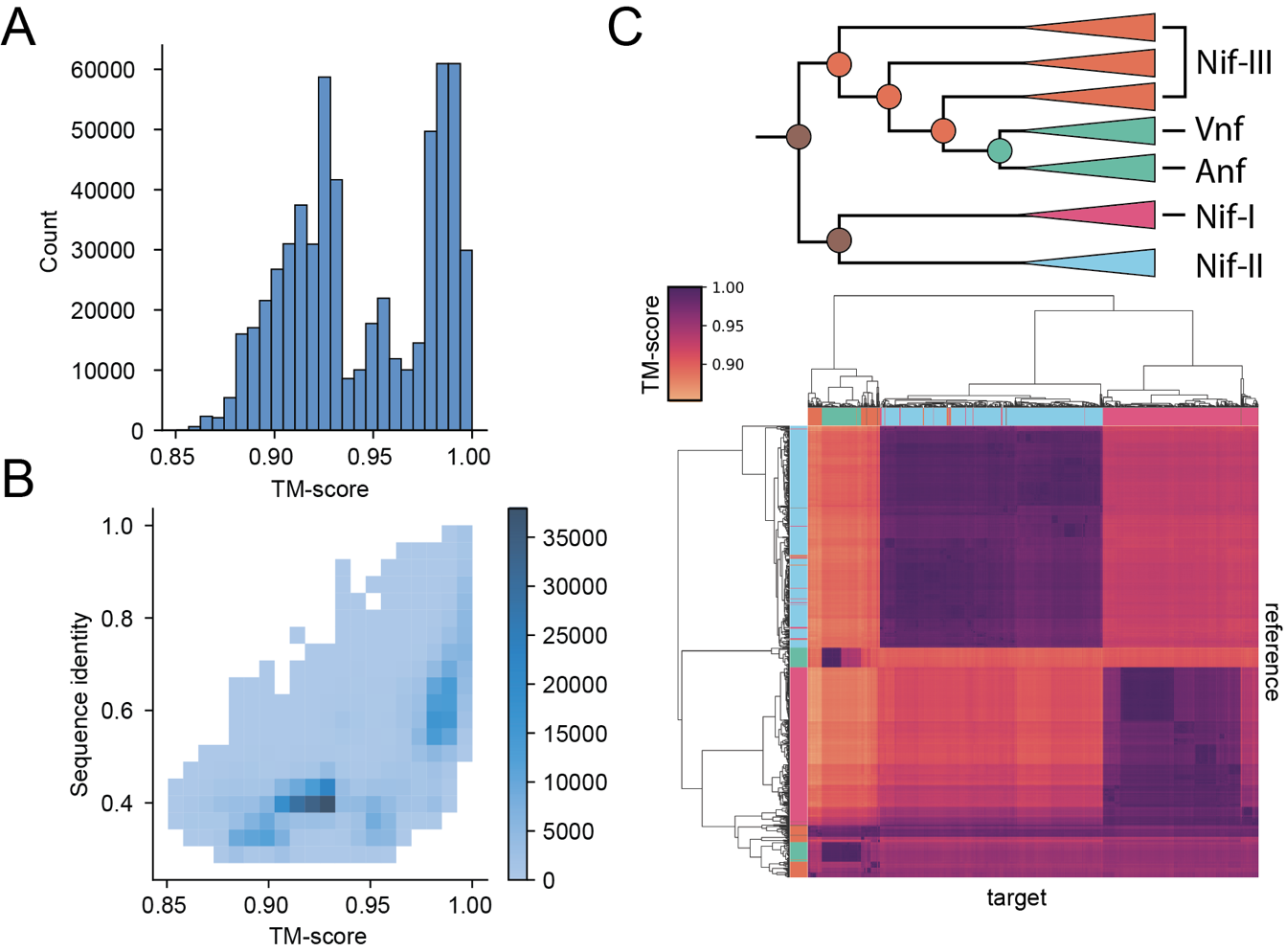

**Figure S4**: **DDKK TM-score relationships. A)** TM-score histogram. TM-score can range from 0 (no similarity) to 1 (identical). The lowest TM-score in this histogram is above 0.85. **B)** Bi-dimensional histogram relating TM-score and sequence identity. **C)** Hierarchical clustering of predicted nitrogenase structures based on structural similarity (TM-score). The colors on the heatmap correspond to the phylogenetic groups represented on **Figure 2C**.

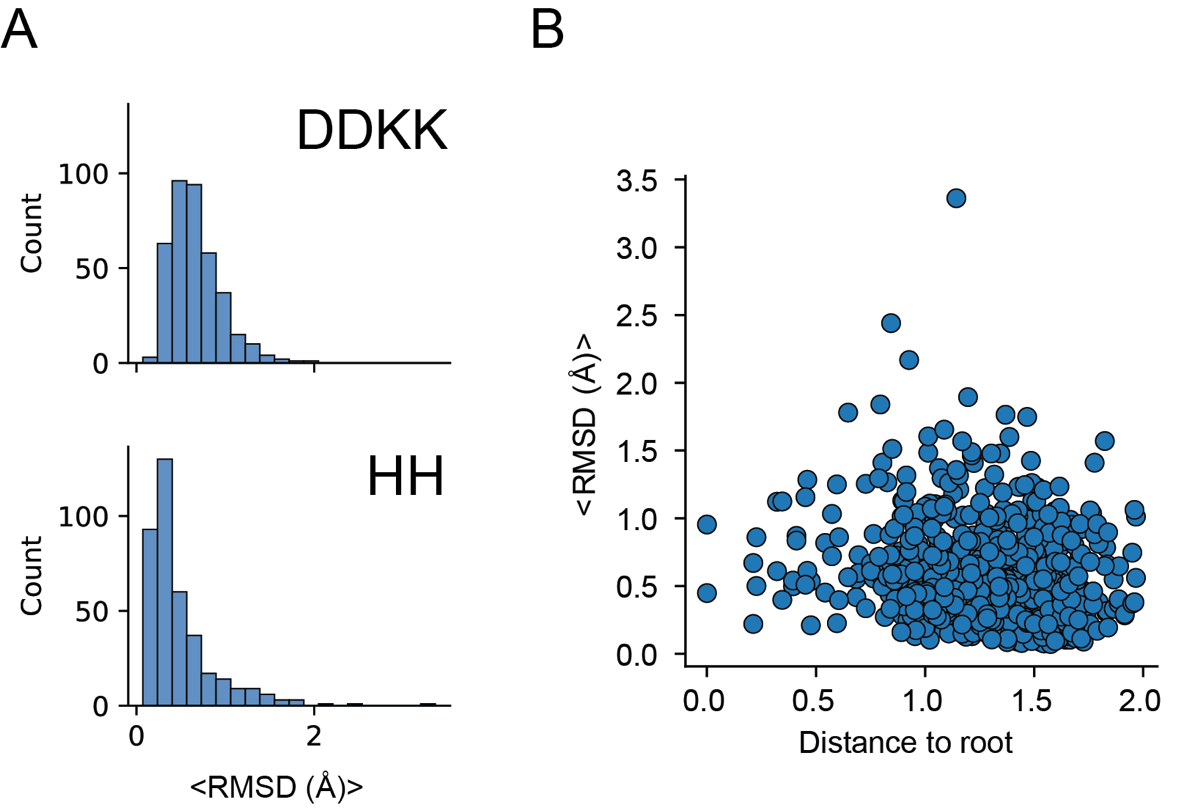

**Figure S5: Structural variation found among sequence variants of ancestral nodes.** Structural similarity is measured as the average RMSD for the backbone alpha-C-atoms of all the alternative (*“alt”*) variants against the maximum-likelihood ancestral variant. **A)** Histogram of the RMSD (Å) for both component 1 (DDKK structures) and 2 (HH structures). **B)** Distance to root versus RMSD (Å) for each ancestral node.

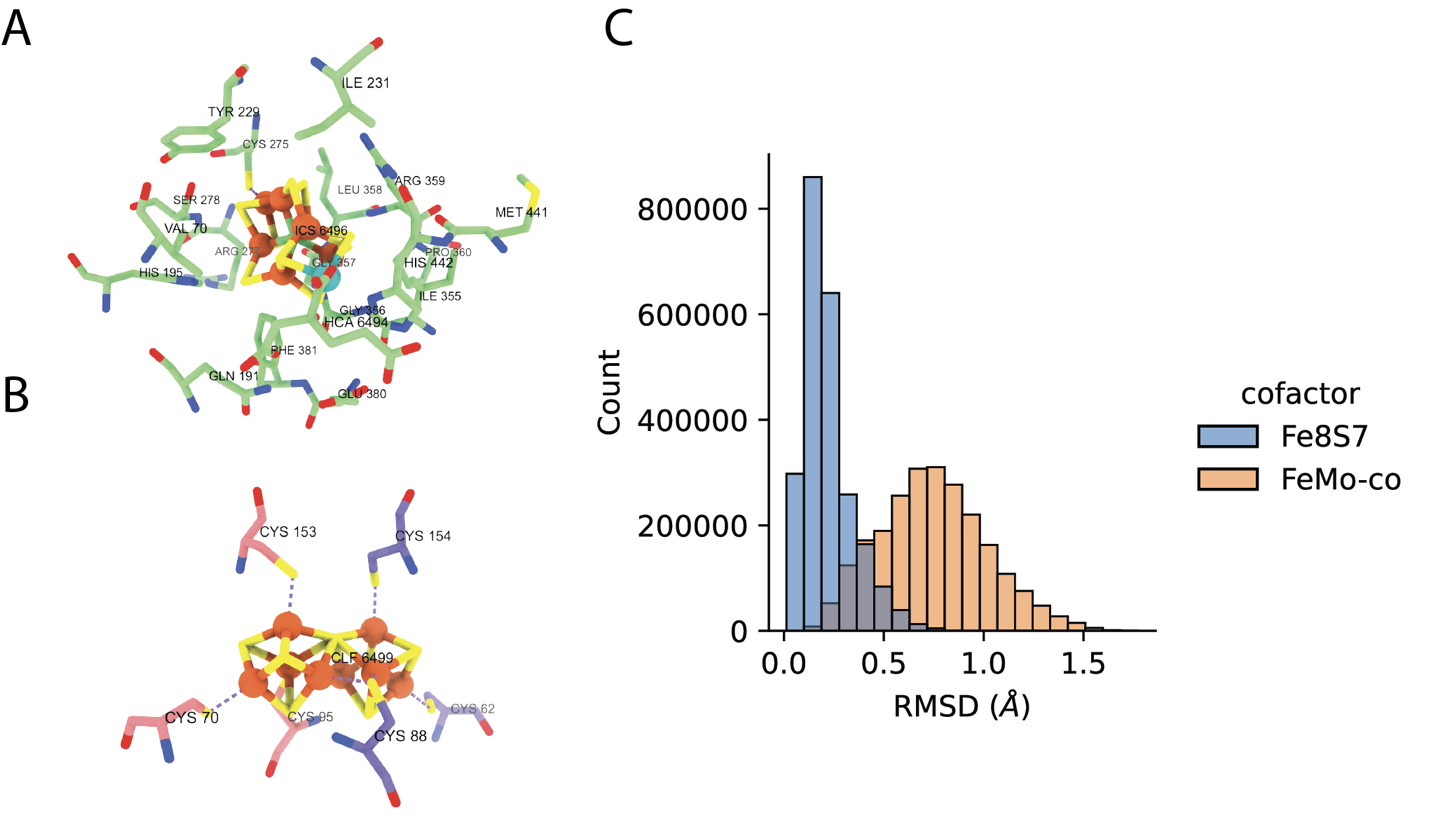

**Figure S6**: **Active site RMSD.** **A)** Selected residues around the M-site (FeMo-co) metal cofactor. Structural snapshot obtained from 3U7Q (Nif *Azotobacter vinelandii*). **B)** Selected residues around the P-site (Fe8S7) metal cofactor. Structural snapshot obtained again from 3U7Q. Residues highlighted in pink correspond to the D subunit, while those in purple correspond to the K subunit. **C)** Alpha-Carbon RMSD for the selected residues in panels A and B.

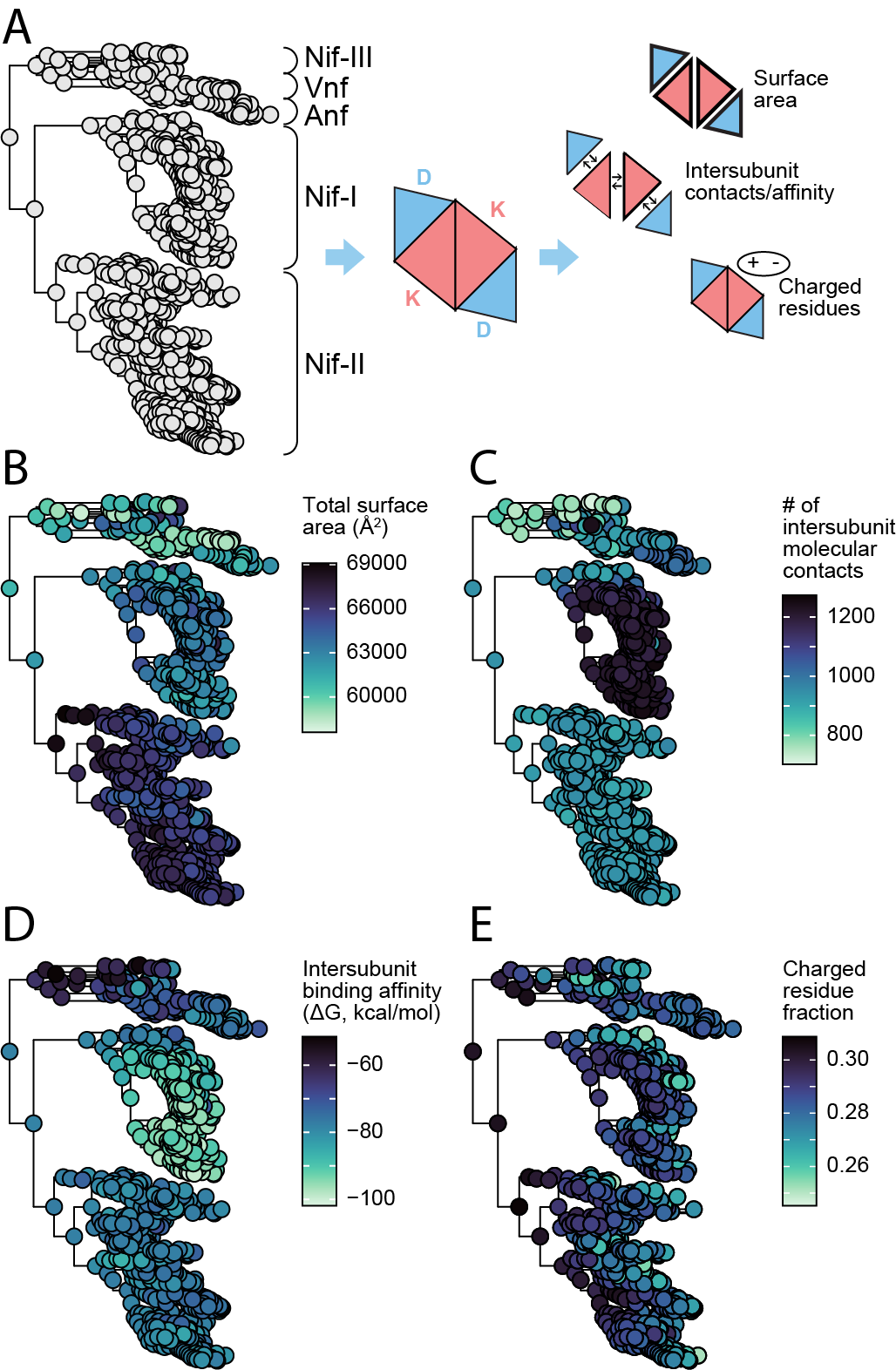

**Figure S7: Phylogenetic analysis of structural features in nitrogenase DDKK proteins.** **A)** Overview of structural attributes calculated from predicted nitrogenase structures. **B-G)** Structural attributes of extant and ancestral nitrogenases mapped to the nitrogenase phylogeny in A).

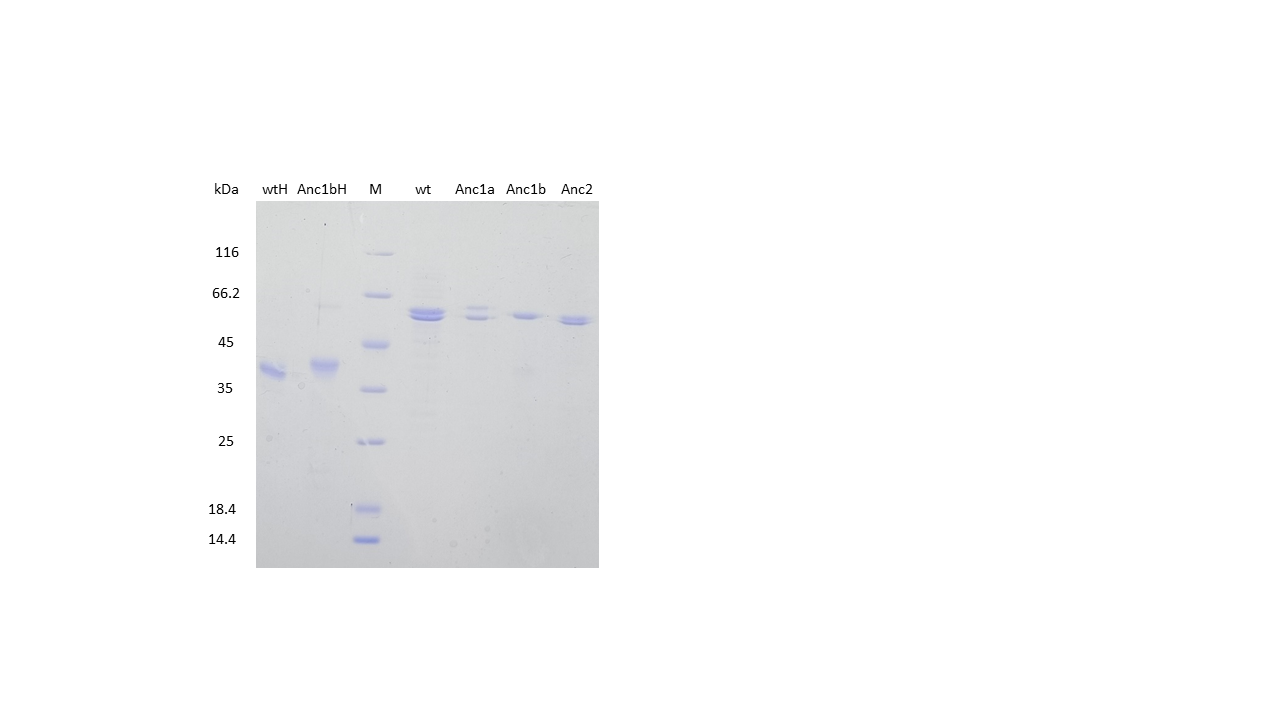

**Figure S8**: SDS-PAGE of the purified proteins. WT H and DK from A. vinelandii are shown for comparison. M indicates marker.

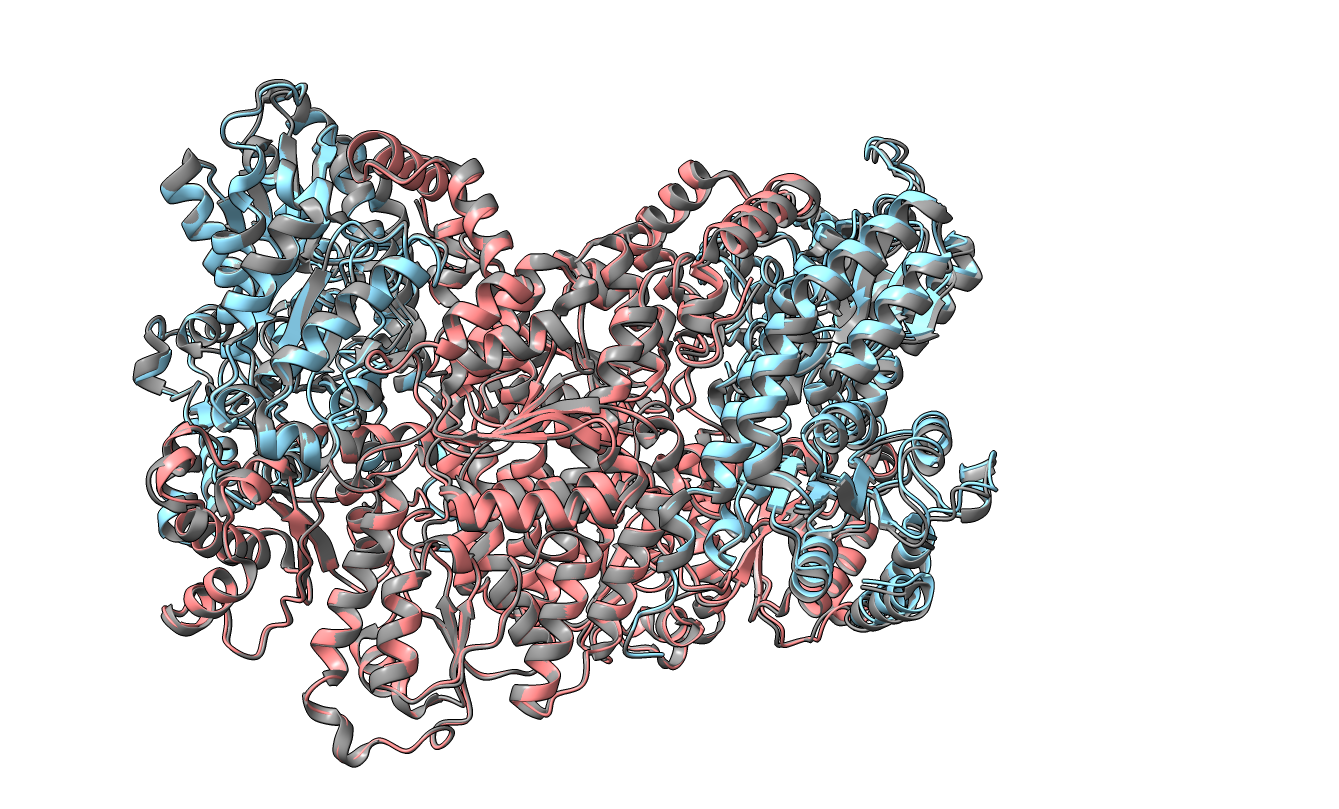

**Figure S9**: Polypeptide chain of WT (colored grey) and Anc1B (colored blue and red) are aligned on the left D subunit (in blue), showing the shift of twofold symmetry axis relating the two DK heterotetramer for the right half of the heterotetramer.

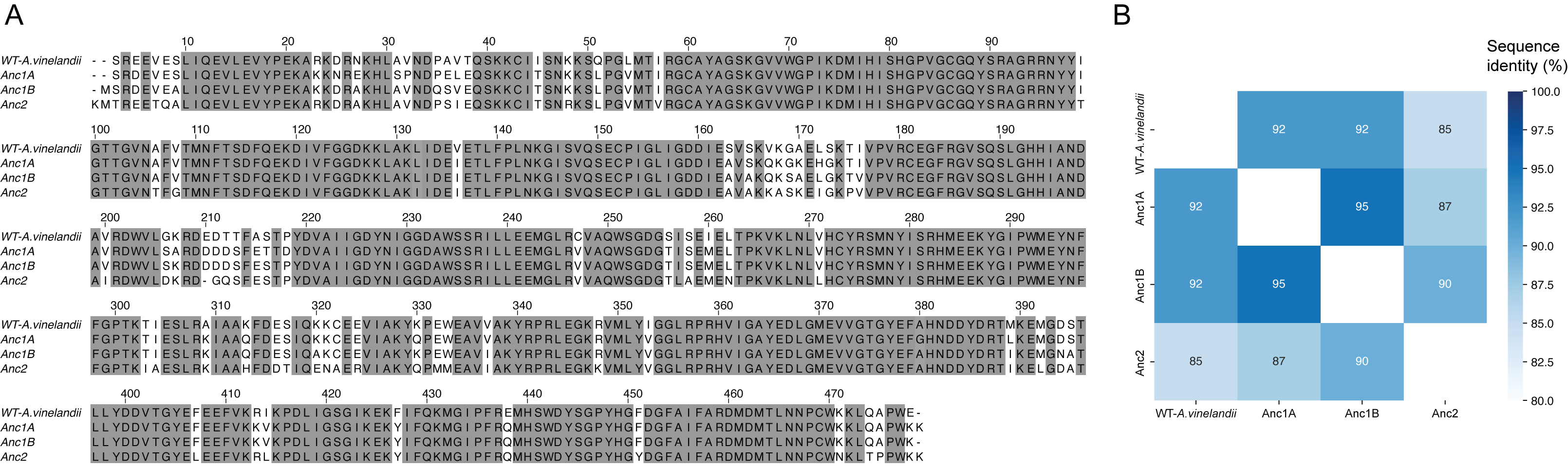

**Figure S10: A)** Sequence alignment for D-subunit from crystallized ancestral nitrogenases. **B)** Sequence identity matrix for D-subunit of crystallized ancestral nitrogenases. All other sequence characteristics of ancestral nitrogenase “Anc” variants are available at Garcia et al., 2022.

**Table S1.** Crystallographic or cryo-EM nitrogenase structures presently available in the Protein Data Bank (<https://www.rcsb.org/>), accessed September 2023. H,D,K and G refer to the respective H, D, K and G-subunits of nitrogenase. The numbering refers to the oligomeric state for each of the subunit in the complex.

| **PDB ID** | **Organism** | Resolution | Year | Type | Stoichiometry |
| --- | --- | --- | --- | --- | --- |
| 8BOQ | *Azotobacter vinelandii* | 1.55 | 2023 | Anf | D2G2K2 |
| 8DFD | *Azotobacter vinelandii* | 2.12 | 2023 | Nif | D2H4K2 |
| 8DBY | *Azotobacter vinelandii* | 2.26 | 2023 | Nif | D2K2 |
| 8DFC | *Azotobacter vinelandii* | 2.48 | 2023 | Nif | D2H2K2 |
| 1NIP | *Azotobacter vinelandii* | 2.90 | 1992 | Nif | H2 |
| 2MIN | *Azotobacter vinelandii* | 2.03 | 1997 | Nif | D2K2 |
| 3MIN | *Azotobacter vinelandii* | 2.03 | 1997 | Nif | D2K2 |
| 1N2C | *Azotobacter vinelandii* | 3.00 | 1997 | Nif | D2H4K2 |
| 2NIP | *Azotobacter vinelandii* | 2.20 | 1998 | Nif | H2 |
| 1FP6 | *Azotobacter vinelandii* | 2.15 | 2000 | Nif | H4 |
| 1DE0 | *Azotobacter vinelandii* | 2.40 | 2000 | Nif | H2 |
| 1G5P | *Azotobacter vinelandii* | 2.20 | 2001 | Nif | H2 |
| 1G20 | *Azotobacter vinelandii* | 2.20 | 2001 | Nif | D2H4K2 |
| 1G1M | *Azotobacter vinelandii* | 2.25 | 2001 | Nif | H2 |
| 1FP4 | *Azotobacter vinelandii* | 2.50 | 2001 | Nif | D2K2 |
| 1G21 | *Azotobacter vinelandii* | 3.00 | 2001 | Nif | D2H4K2 |
| 1M1N | *Azotobacter vinelandii* | 1.16 | 2002 | Nif | D4K4 |
| 1L5H | *Azotobacter vinelandii* | 2.30 | 2002 | Nif | D1K1 |
| 1M34 | *Azotobacter vinelandii* | 2.30 | 2002 | Nif | D4H8K4 |
| 1M1Y | *Azotobacter vinelandii* | 3.20 | 2002 | Nif | D4H8K4 |
| 1RW4 | *Azotobacter vinelandii* | 2.50 | 2004 | Nif | H1 |
| 1XD8 | *Azotobacter vinelandii* | 2.70 | 2004 | Nif | H2 |
| 1XD9 | *Azotobacter vinelandii* | 2.80 | 2004 | Nif | H2 |
| 1XDB | *Azotobacter vinelandii* | 2.80 | 2004 | Nif | H2 |
| 1XCP | *Azotobacter vinelandii* | 3.20 | 2004 | Nif | H4 |
| 2AFH | *Azotobacter vinelandii* | 2.10 | 2005 | Nif | D2H2K2 |
| 4WZB | *Azotobacter vinelandii* | 2.30 | 2005 | Nif | D2H4K2 |
| 2AFI | *Azotobacter vinelandii* | 3.10 | 2005 | Nif | D4H8K4 |
| 2C8V | *Azotobacter vinelandii* | 2.50 | 2006 | Nif | H1 |
| 3K1A | *Azotobacter vinelandii* | 2.23 | 2010 | Nif | D2K2 |
| 3U7Q | *Azotobacter vinelandii* | 1.00 | 2011 | Nif | D2K2 |
| 4TKU | *Azotobacter vinelandii* | 1.43 | 2014 | Nif | D2K2 |
| 4TKV | *Azotobacter vinelandii* | 1.50 | 2014 | Nif | D2K2 |
| 4ND8 | *Azotobacter vinelandii* | 2.00 | 2014 | Nif | D2K2 |
| 5BVH | *Azotobacter vinelandii* | 1.53 | 2015 | Nif | D2K2 |
| 5BVG | *Azotobacter vinelandii* | 1.60 | 2015 | Nif | D2K2 |
| 5CX1 | *Azotobacter vinelandii* | 1.75 | 2015 | Nif | D8K8 |
| 4WZA | *Azotobacter vinelandii* | 1.90 | 2015 | Nif | D2H4K2 |
| 4XPI | *Azotobacter vinelandii* | 1.97 | 2015 | Nif | D2K2 |
| 4WNA | *Azotobacter vinelandii* | 2.00 | 2015 | Nif | D2K2 |
| 6BBL | *Azotobacter vinelandii* | 1.68 | 2017 | Nif | D2K2 |
| 5VQ4 | *Azotobacter vinelandii* | 2.30 | 2017 | Nif | D2K2 |
| 6CDK | *Azotobacter vinelandii* | 2.10 | 2018 | Nif | D2K2 |
| 6N4L | *Azotobacter vinelandii* | 1.13 | 2019 | Nif | H1 |
| 6O7M | *Azotobacter vinelandii* | 1.40 | 2019 | Nif | D2K2 |
| 6N4M | *Azotobacter vinelandii* | 1.58 | 2019 | Nif | H1 |
| 6O0B | *Azotobacter vinelandii* | 1.60 | 2019 | Nif | H2 |
| 6OP3 | *Azotobacter vinelandii* | 1.60 | 2019 | Nif | D2K2 |
| 6O7P | *Azotobacter vinelandii* | 1.70 | 2019 | Nif | D2K2 |
| 6OP1 | *Azotobacter vinelandii* | 1.70 | 2019 | Nif | D2K2 |
| 6O7N | *Azotobacter vinelandii* | 1.75 | 2019 | Nif | D2K2 |
| 6N4K | *Azotobacter vinelandii* | 1.76 | 2019 | Nif | H2 |
| 6O7O | *Azotobacter vinelandii* | 1.89 | 2019 | Nif | D2K2 |
| 6OP2 | *Azotobacter vinelandii* | 1.90 | 2019 | Nif | D2K2 |
| 6N4J | *Azotobacter vinelandii* | 1.95 | 2019 | Nif | H2 |
| 6O7Q | *Azotobacter vinelandii* | 2.00 | 2019 | Nif | D2K2 |
| 6O7L | *Azotobacter vinelandii* | 2.26 | 2019 | Nif | D2K2 |
| 6O7S | *Azotobacter vinelandii* | 2.27 | 2019 | Nif | D2K2 |
| 6O7R | *Azotobacter vinelandii* | 2.27 | 2019 | Nif | D2K2 |
| 6OP4 | *Azotobacter vinelandii* | 2.30 | 2019 | Nif | D2K2 |
| 6VXT | *Azotobacter vinelandii* | 1.74 | 2020 | Nif | D2K2 |
| 6UG0 | *Azotobacter vinelandii* | 1.83 | 2020 | Nif | D2K2 |
| 7JRF | *Azotobacter vinelandii* | 1.33 | 2021 | Nif | D2K2 |
| 7TPW | *Azotobacter vinelandii* | 1.18 | 2022 | Nif | H1 |
| 7TPO | *Azotobacter vinelandii* | 1.35 | 2022 | Nif | H1 |
| 7TPX | *Azotobacter vinelandii* | 1.35 | 2022 | Nif | H1 |
| 7TPN | *Azotobacter vinelandii* | 1.38 | 2022 | Nif | H1 |
| 7TNE | *Azotobacter vinelandii* | 1.39 | 2022 | Nif | H1 |
| 7TQF | *Azotobacter vinelandii* | 1.45 | 2022 | Nif | H1 |
| 7TQI | *Azotobacter vinelandii* | 1.47 | 2022 | Nif | H1 |
| 7TPY | *Azotobacter vinelandii* | 1.48 | 2022 | Nif | H1 |
| 7TQJ | *Azotobacter vinelandii* | 1.48 | 2022 | Nif | H1 |
| 7TQK | *Azotobacter vinelandii* | 1.48 | 2022 | Nif | H1 |
| 7TPV | *Azotobacter vinelandii* | 1.49 | 2022 | Nif | H1 |
| 7TQH | *Azotobacter vinelandii* | 1.49 | 2022 | Nif | H1 |
| 7T4H | *Azotobacter vinelandii* | 1.51 | 2022 | Nif | H1 |
| 7TQC | *Azotobacter vinelandii* | 1.53 | 2022 | Nif | H1 |
| 7TQE | *Azotobacter vinelandii* | 1.59 | 2022 | Nif | H1 |
| 7TQ9 | *Azotobacter vinelandii* | 1.60 | 2022 | Nif | H1 |
| 7TPZ | *Azotobacter vinelandii* | 1.71 | 2022 | Nif | H1 |
| 7TQ0 | *Azotobacter vinelandii* | 1.81 | 2022 | Nif | H1 |
| 8DBX | *Azotobacter vinelandii* | 1.92 | 2023 | Nif | D2K2 |
| 8CRS | *Azotobacter vinelandii* | 2.04 | 2023 | Nif | D2K2 |
| 8ENM | *Azotobacter vinelandii* | 2.14 | 2023 | Nif | D2K2 |
| 8ENL | *Azotobacter vinelandii* | 2.37 | 2023 | Nif | D2K2 |
| 8BTS | *Azotobacter vinelandii* | 3.03 | 2023 | Nif | D4K4 |
| 5N6Y | *Azotobacter vinelandii* | 1.35 | 2017 | Vnf | D2G2K2 |
| 6FEA | *Azotobacter vinelandii* | 1.20 | 2018 | Vnf | D2G2K2 |
| 6Q93 | *Azotobacter vinelandii* | 2.20 | 2018 | Vnf | H8 |
| 7ADR | *Azotobacter vinelandii* | 1.00 | 2020 | Vnf | D2G2K2 |
| 7ADY | *Azotobacter vinelandii* | 1.05 | 2020 | Vnf | D2G2K2 |
| 7AIZ | *Azotobacter vinelandii* | 1.05 | 2021 | Vnf | D2G2K2 |
| 1MIO | *Clostridium pasteurianum* | 3.00 | 1993 | Nif | D2K2 |
| 1CP2 | *Clostridium pasteurianum* | 1.93 | 1998 | Nif | H2 |
| 4WN9 | *Clostridium pasteurianum* | 1.90 | 2015 | Nif | D2K2 |
| 5VQ3 | *Clostridium pasteurianum* | 1.72 | 2017 | Nif | D2K2 |
| 5VPW | *Clostridium pasteurianum* | 1.85 | 2017 | Nif | D2K2 |
| 5KOH | *Gluconacetobacter*  *diazotrophicus* PA1 5 | 1.83 | 2016 | Nif | D2K2 |
| 5KOJ | *Gluconacetobacter*  *diazotrophicus* PA1 5 | 2.59 | 2016 | Nif | D2K2 |
| 1QH8 | *Klebsiella pneumoniae* | 1.60 | 1999 | Nif | D2K2 |
| 1QH1 | *Klebsiella pneumoniae* | 1.60 | 1999 | Nif | D2K2 |
| 1QGU | *Klebsiella pneumoniae* | 1.60 | 1999 | Nif | D2K2 |
| 1H1L | *Klebsiella pneumoniae* | 1.90 | 2002 | Nif | D2K2 |
| 6NZJ | *Methanosarcina*  *acetivorans* C2A | 2.40 | 2019 | Nif | H2 |

**Table S2**: Data collection and refinement statistics for Anc2

| **Data collection** |  |
| --- | --- |
| Wavelength (Å) | 1.000 |
| Spherical resolution (Å) | 110.49-1.816 (2.013-1.816) |
| Limiting resolution (Å) along |  |
| a* | 2.417 |
| b* | 1.984 |
| c* | 1.816 |
| Space group | P 2_1_2_1_2_1_ |
| Unit cell | 74.936, 130.133, 209.137, 90, 90, 90, |
| Total reflections | 1683610 (81981) |
| Unique reflections | 121997 (6101) |
| Multiplicity | 13.80 (13.4) |
| Completeness, spherical (%) | 66.51 (12.6) |
| Completeness, ellipsoidal (%) | 95.0 (70.2) |
| Mean I/sigma(I) | 10.0 (1.6) |
| Wilson B-factor | 21.85 |
| R-merge (Weiss and Hilgenfeld,1997) | 0.202 (1.696) |
| R-meas | 0.210 (1.772) |
| R-pim (Weiss and Hilgenfeld, 1997) | 0.056 (0.478) |
| CC_1/2_ (Karplus and Diederichs, 2012) | 0.996 (0.698) |
| **Refinement** |  |
| R-work | 0.1864 |
| R-free | 0.2305 |
| RMS(bonds) | 0.024 |
| RMS(angles) | 1.68 |
| Ramachandran favored (%) | 96.44 |
| Ramachandran allowed (%) | 3.36 |
| Ramachandran outliers (%) | 0.20 |
| Rotamer outliers (%) | 1.62 |
| Clashscore | 3.88 |
| Average B-factor | 27.72 |
| ..macromolecules | 27.57 |
| ..ligands | 23.48 |
| ..solvent | 29.97 |

Statistics for the highest-resolution shell are shown in parentheses.

**Table S3:** Data collection and refinement statistics for Anc1A

| **Data collection** |  |
| --- | --- |
| Wavelength (Å) | 0.8856 |
| Spherical resolution (Å) | 19.791-2.658 (2.912-2.658) |
| Limiting resolution (Å) along |  |
| a* | 2.934 |
| b* | 2.656 |
| c* | 2.913 |
| Space group | P 2_1_2_1_2_1_ |
| Unit cell | 70.29, 138.61, 208.58, 90.0, 90.0, 90.0 |
| Total reflections | 595126 (29588) |
| Unique reflections | 46401 (2321) |
| Multiplicity | 12.8 (12.7) |
| Completeness, spherical (%) | 78.07 (16.5) |
| Completeness, ellipsoidal (%) | 93.7 (51.8) |
| Mean I/sigma(I) | 7.1 (1.4) |
| Wilson B-factor | 39.22 |
| R-merge (Weiss and Hilgenfeld,1997) | 0.368 (2.046) |
| R-meas | 0.384 (2.131) |
| R-pim (Weiss and Hilgenfeld,1997) | 0.106 (0.593) |
| CC_1/2_ (Karplus and Diederichs, 2012) | 0.989 (0.425) |
| **Refinement** |  |
| R-work | 0.1796 |
| R-free | 0.2220 |
| RMS(bonds) | 0.022 |
| RMS(angles) | 1.44 |
| Ramachandran favored (%) | 95.53 |
| Ramachandran allowed (%) | 4.47 |
| Ramachandran outliers (%) | 0.00 |
| Rotamer outliers (%) | 5.57 |
| Clashscore | 3.95 |
| Average B-factor | 42.75 |
| ..macromolecules | 42.91 |
| ..ligands | 42.28 |
| solvent | 23.71 |

Statistics for the highest-resolution shell are shown in parentheses.

**Table S4**: Data collection and refinement statistics for Anc1B HH

| **Data collection** |  |
| --- | --- |
| Wavelength (Å) | 0.8856 |
| Spherical resolution (Å) | 69.75 - 2.434 (2.521 - 2.434) |
| Limiting resolution (Å) along |  |
| a* | 2.434 |
| b* | 2.468 |
| c* | 2.828 |
| Space group | C 12_1_ |
| Unit cell | 123.35, 109.78, 95.21, 90.0, 118.6, 90.0 |
| Total reflections | 45985 (2294) |
| Unique reflections | 26823 (1341) |
| Multiplicity | 1.7 (1.7) |
| Mean I/sigma(I) | 2.74 (1.52) |
| Wilson B-factor | 28.93 |
| R-merge (Weiss and Hilgenfeld,1997) | 0.096 (0.321) |
| R-meas | 0.132 (0.447) |
| R-pim (Weiss and Hilgenfeld,1997) | 0.09 (0.310) |
| CC_1/2_ (Karplus and Diederichs, 2012) | 0.98 (0.716) |
| **Refinement** |  |
| R-work | 0.2023 |
| R-free | 0.2722 |
| RMS(bonds) | 0.008 |
| RMS(angles) | 1.49 |
| Ramachandran favored (%) | 95.72 |
| Ramachandran allowed (%) | 3.75 |
| Ramachandran outliers (%) | 0.52 |
| Rotamer outliers (%) | 9.31 |
| Clashscore | 7.19 |
| Average B-factor | 29.54 |
| macromolecules | 29.67 |
| ligands | 19.91 |
| solvent | 16.18 |

Statistics for the highest-resolution shell are shown in parentheses.

**Table S5:** Data collection and refinement statistics for Anc1B DDKK

| **Data collection** |  |
| --- | --- |
| Wavelength (Å) | 0.8856 |
| Spherical resolution (Å) | 109.828-1.824 (2.105-1.824) |
| Limiting resolution (Å) along |  |
| a* | 2.805 |
| b* | 2.321 |
| c* | 1.821 |
| Space group | P 2_1_2_1_2_1_ |
| Unit cell | 77.08, 128.98, 209.45, 90.0, 90.0, 90.0 |
| Total reflections | 1198687 (60971) |
| Unique reflections | 91363 (4568) |
| Multiplicity | 13.1 (13.1) |
| Completeness, spherical (%) | 49.1 (7.1) |
| Completeness, ellipsoidal (%) | 94.4 (77.0) |
| Mean I/sigma(I) | 7.5 (2.0) |
| Wilson B-factor | 20.64 |
| R-merge (Weiss and Hilgenfeld,1997) | 0.237 (1.419) |
| R-meas | 0.247 (1.476) |
| R-pim (Weiss and Hilgenfeld,1997) | 0.068 (0.401) |
| CC_1/2_ (Karplus and Diederichs, 2012) | 0.997 (0.727) |
| **Refinement** |  |
| R-work | 0.2063 |
| R-free | 0.2686 |
| RMS(bonds) | 0.075 |
| RMS(angles) | 1.6982 |
| Ramachandran favored (%) | 95.78 |
| Ramachandran allowed (%) | 3.97 |
| Ramachandran outliers (%) | 0.25 |
| Rotamer outliers (%) | 2.46 |
| Clashscore | 6.90 |
| Average B-factor | 25.55 |
| macromolecules | 25.77 |
| ligands | 18.94 |
| solvent | 21.33 |
| Number of TLS groups | 4 |

Statistics for the highest-resolution shell are shown in parentheses.

**Table S6:** Structural features analyzed in the massive nitrogenase structure prediction, with their respective programs.

| Variable | Program | Explanation |
| --- | --- | --- |
| Total area | FreeSASA  (Mitternacht, 2016) | Exposed protein surface, in Å2 |
| Buried area | FreeSASA (Mitternacht 2016) | The sum of the individual chains surfaces minus the protein surface. |
| Polar area | FreeSASA (Mitternacht 2016) |  |
| Polar buried area | FreeSASA (Mitternacht 2016) |  |
| Apolar area | FreeSASA (Mitternacht 2016) |  |
| Apolar buried area | FreeSASA (Mitternacht 2016) |  |
| Intermolecular contacts | Prodigy (Vangone and Bonvin, 2015) | Number of contacts between residues of different chains. |
| Charged-charged contacts | Prodigy (Vangone and Bonvin 2015) | Number of interactions between charged residues of different chains. |
| Charged-polar contacts | Prodigy (Vangone and Bonvin 2015) | Number of interactions between charged residues and polar residues of different chains. |
| Charged-apolar contacts | Prodigy (Vangone and Bonvin 2015) | Number of interactions between charged residues and apolar residues of different chains. |
| Apolar-polar contacts | Prodigy (Vangone and Bonvin 2015) | Number of interactions between apolar residues and polar residues of different chains. |
| Apolar Non-Interacting-Surface | Prodigy (Vangone and Bonvin 2015) | See (Kastritis et al., 2014) |
| Charged Non-Interacting-Surface | Prodigy (Vangone and Bonvin 2015) | See (Kastritis et al. 2014) |
| Binding affinity | Prodigy (Vangone and Bonvin 2015) | Free energy of binding of the complex, in kcal/mol. The lower, the more stable the complex. |
| Gaussian Normal Mode eigenvalues (first five) | Prody (Bakan et al., 2011; Zhang et al., 2021) | Lowest eigenvalues associated with the Gaussian Normal Mode analysis. These values are associated with the frequency of normal mode oscillations (Bauer et al., 2019) |
| Radius of gyration | Prody (Bakan et al. 2011; Zhang et al. 2021) | See (Lobanov et al., 2008) |
| Residue-Residue average shortest path | RING (Clementel et al., 2022) /NetworkX (Hagberg et al., 2008) | Mean length of the shortest path joining two random nodes in a connected network. |
| Residue average clustering | RING (Clementel et al. 2022) /NetworkX (Hagberg et al. 2008) | Number of edges connecting the node and its neighbors between them divided by the number of edges of a complete subgraph involving all its neighbors. |
| Residue network density | RING (Clementel et al. 2022) /NetworkX (Hagberg et al. 2008) | Number of edges divided the number of edges that would be expected if the graph was complete. |
| Residue average degree | RING (Clementel et al. 2022) /NetworkX (Hagberg et al. 2008) | Mean number of neighbors of each residue. |
